# Methylphenidate as a causal test of translational and basic neural coding hypotheses

**DOI:** 10.1101/2021.09.12.459995

**Authors:** Amy M. Ni, Brittany S. Bowes, Douglas A. Ruff, Marlene R. Cohen

## Abstract

Most systems neuroscience studies fall into one of two categories: basic science work aimed at understanding the relationship between neurons and behavior, or translational work aimed at developing treatments for neuropsychiatric disorders. Here we use these two approaches to inform and enhance each other. Our study both tests hypotheses about basic science neural coding principles and elucidates the neuronal mechanisms underlying new, clinically relevant behavioral effects of systemically administered methylphenidate (Ritalin). We discovered that orally administered methylphenidate, used clinically to treat Attention Deficit Hyperactivity Disorder (ADHD) and generally to enhance cognition (Lakhan & Kirchgessner, 2012; Maher, 2008), increases spatially selective visual attention, enhancing visual performance at only the attended location. And as predicted by our previous work (Ni et al., 2018), we found that this causal manipulation enhances vision in rhesus macaques specifically when it decreases the mean correlated variability of neurons in visual area V4. Our findings demonstrate that the visual system is a platform for understanding the neural underpinnings of both complex cognitive processes (basic science) and neuropsychiatric disorders (translation). Addressing basic science hypotheses, our results are consistent with a scenario in which methylphenidate has cognitively specific effects by working through naturally selective cognitive mechanisms. Clinically, our findings suggest that the often staggeringly specific symptoms of neuropsychiatric disorders may be caused and treated by leveraging general mechanisms.

## INTRODUCTION

Studying the behavioral and neuronal effects of stimulants such as methylphenidate is important for both translational and basic science reasons. It is of translational importance because stimulants are widely used by adults and children but their neuronal mechanisms remain unclear (Mueller et al., 2017). More than 6% of children in the United States are prescribed stimulants to treat ADHD (Visser et al., 2014). Additionally, one fifth of polled Nature readers report using these stimulants without prescription to enhance performance (Maher, 2008), with this number thought to be much larger among college students (Lakhan & Kirchgessner, 2012). These stimulants are frequently used both with and without prescription with the intention of improving selective attention, which allows one to focus on a desired target and tune out distractors (Maunsell, 2015). However, despite the frequent goal of achieving selective changes in performance, most behavioral and neuroscientific studies of stimulants have focused on examining overall performance changes related to global processes such as motivation and vigilance (Bagot & Kaminer, 2014; Koelega, 1993; Lakhan & Kirchgessner, 2012; McLellan et al., 2016; Mueller et al., 2017; Murray, 2010; Pietrzak et al., 2006; Spencer, et al., 2013; Swanson et al., 2011; Wickens et al., 2011).

Studying stimulants is also important because it provides a strong, causal test of basic science hypotheses about how groups of neurons affect visually guided behaviors. In a previous study (Ni et al., 2018), we demonstrated that there is a robust relationship between the magnitude of correlated variability in visual cortex (the shared trial-to-trial variability of pairs of neurons in response to repeated presentations of the same stimulus; Cohen & Kohn, 2011) and the ability of rhesus monkeys to detect changes in the orientation of a visual stimulus. This relationship between neuronal populations in visual area V4 and performance persisted whether correlated variability and behavior were changed by spatial attention on fast timescales, perceptual learning over several weeks, or factors outside experimenter control. These observations led to the hypothesis that a cognitive process, neuropsychiatric disorder, or causalmanipulation should affect performance on this task precisely when it affects correlated variability in V4. Methylphenidate as a causal manipulation comprises a strong test of this hypothesis because it has widespread effects on the dopamine system throughout the brain (Arnsten, 2006; Noudoost & Moore, 2011b), and it is unknown whether a systemically administered stimulant can have such specific effects on neuronal activity.

## RESULTS

To test our basic science hypotheses and investigate the clinically relevant behavioral and neuronal effects of methylphenidate, we administered methylphenidate and recorded populations of V4 neurons in rhesus monkeys trained to perform a perceptually challenging visual task with a spatial attention component. We chose oral administration because this is the most common means of methylphenidate administration (Pietrzak et al., 2006) and to test the effects of a systemic manipulation of the attentional system on the activity of a neuronal population in sensory cerebral cortex.

On alternating days, a monkey drank either sugar water with methylphenidate mixed in or a placebo of only sugar water (Soto et al., 2012). The sugar water with or without methylphenidate was administered 30 minutes prior to behavioral testing (Gamo et al., 2010).

The heart of our analysis approach is to compare pairs of experimental sessions with matched stimulus and task parameters (see **Methods**) that were conducted on adjacent days. Each pair of sessions included one in which we administered methylphenidate and one in which we administered a placebo control.

We used between 2-6 mg/kg (see **Methods**; Kodama et al., 2017; Oemisch et al., 2016; Rajala et al., 2012; 2015; 2020), and the data from all dosages were included together in the analyses to avoid best dose analyses (Soto et al., 2013; while our goal was to use the systemic administration of methylphenidate as a causal test of our hypotheses, not to test for dose-dependent effects, we have included analyses per dosage in the **Supplementary Figures**).

To measure the effects of methylphenidate on selective attention, we trained three rhesus monkeys to perform the visual change-detection task that we used to manipulate spatial attention in our previous work (**Fig. 1a**; Cohen & Maunsell, 2009; Ni et al., 2018). The monkey fixated a central point while two peripheral Gabor stimuli flashed on and off. At a random and unsignaled time, the orientation of one stimulus changed slightly. The monkey was rewarded for making an eye movement toward the changed stimulus. We manipulated spatial attention using a classic Posner spatial attention paradigm (Posner 1980): before each block of trials, the monkey was cued to attend to the location where the orientation change was most likely to happen. The orientation change occurred at the attended location 80% of the time, and the animal was rewarded for detecting changes at both the attended and unattended location. The attended location alternated between the left and right locations on each new block of trials.

**Figure 1.**
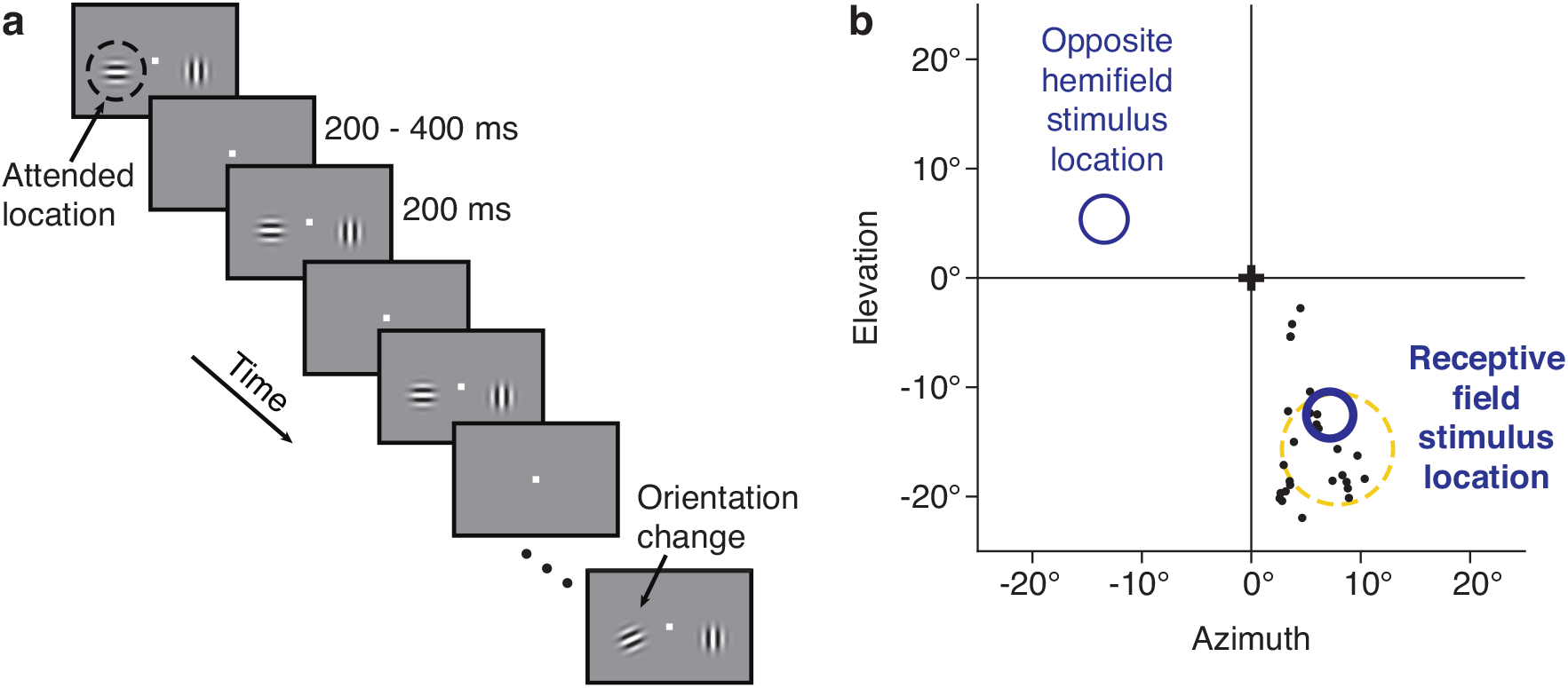
Behavioral and recording methods. (**a**) Orientation change-detection task with a spatial attention manipulation. This task is similar to one we have used in previous studies linking correlated variability in V4 to attention and performance (Cohen & Maunsell, 2009; Ni et al., 2018). The monkey was required to fixate a central spot while two Gabor stimuli flashed on and off, one in the left visual hemifield and one in the right. The monkeys were rewarded for detecting a subtle orientation change that occurred at either the attended location (80% of trials) or the unattended location. The orientation change occurred at a randomized location and time. The attended location was cued using unanalyzed instruction trials at the beginning of each block of trials. The starting orientation of each of the two stimuli was selected randomly per stimulus and per trial from a set of 4-12 orientations. (**b**) Physiological methods. For monkeys 2 and 3, we recorded from chronically implanted microelectrode arrays in visual area V4. We recorded the responses of a few dozen V4 neurons simultaneously. The receptive fields of the recorded neurons typically overlapped both each other and the location of one of the Gabor stimuli (the receptive field stimulus location). The figure depicts, for an example recording session, the centers of the receptive fields of the recorded neurons (black dots), a typical receptive field size and location (dotted yellow circle), and the locations of the two Gabor stimuli (dark blue circles).

For two of the monkeys, we simultaneously recorded the activity of a few dozen neurons in visual area V4 using chronically implanted microelectrode arrays. The two visual stimuli were positioned such that one stimulus overlapped the receptive fields of the recorded V4 neurons (**Fig. 1b**) and the other was in the opposite hemifield.

### Improved motivation

To investigate the many clinically relevant behavioral effects of methylphenidate (Bagot & Kaminer, 2014; Koelega, 1993; Lakhan & Kirchgessner, 2012; Pietrzak et al., 2006; Swanson et al., 2011) in our controlled laboratory setting, we measured many aspects of the monkeys’ behavior and quantitatively compared days on which we administered methylphenidate to their corresponding placebo control days. The most dramatic change was in the amount of time the monkeys engaged in the behavioral task. For our behavioral data sets (see **Methods**), the monkeys controlled the length of the session: the experiment ended when the monkey had not fixated the central spot to initiate a trial for 10 minutes. Even when we matched the total amount of liquid the monkeys received prior to drug and placebo control days to control for any effect of the prior day’s juice intake (**Supp. Fig. 1a**), the monkeys performed the task nearly twice as long on drug than control days (**Fig. 2**). The methylphenidate dosage did not significantly affect working time (**Supp. Fig. 1b**; though see Rajala et al., 2012; 2020).

**Figure 2.**
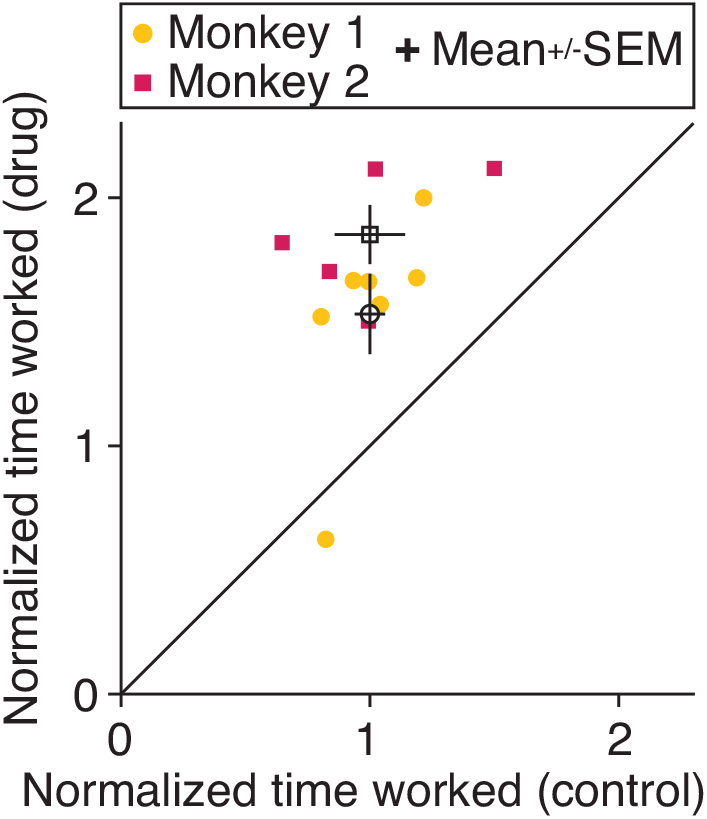
Methylphenidate improves measures of general processes like motivation or work ethic. For a subset of days on which we followed a strict protocol for measuring time engaged on the change-detection task (see **Methods**), the plot depicts the amount of time the monkey engaged in the task each day, normalized to the mean time worked on all placebo control days. Each point is the normalized working time for a drug day (y-axis) and its matched control day (x-axis; adjacent control day with identical stimulus parameters) for each monkey (marker symbols). The open symbols are the mean for each monkey, and error bars represent standard error of the mean (SEM). Both animals worked significantly longer on drug than control days (paired t-tests; Monkey 1: n = 7 pairs of days, t(6) = -4.1, p = 6.1 × 10^-3^; Monkey 2: n = 5 pairs of days, t(4) = -6.6, p = 2.7 × 10^-3^).

### Increased selective attention

Even though we administered methyphenidate systemically, methylphenidate improved behavioral performance on our challenging visual change-detection task at only the attended location (**Fig. 3a**). Methylphenidate did not increase performance at the unattended location (**Fig. 3b**), such that it overall increased the selective effects of attention (the difference in performance between the attended and unattended locations; **Fig. 3c**). Comparing the attention conditions directly demonstrates that the methylphenidate effects were different at the attended versus unattended locations (**Fig. 3c**). The methylphenidate dosage did not significantly affect the animal’s performance on the change-detection task (**Supp. Fig. 2a, b**). There was no indication of a relationship between performance and motivation effects, suggesting distinct mechanisms (**Supp. Fig. 2c, d**). The positive effect of methylphenidate on performance at the attended location was due to both improved visual sensitivity (improving the monkey’s ability to see the difference between the original and changed stimuli in our task; **Supp. Fig. 3a**) and decreased criterion (increasing the readiness of the animal to move its eyes; **Supp. Fig. 3b**).

**Figure 3.**
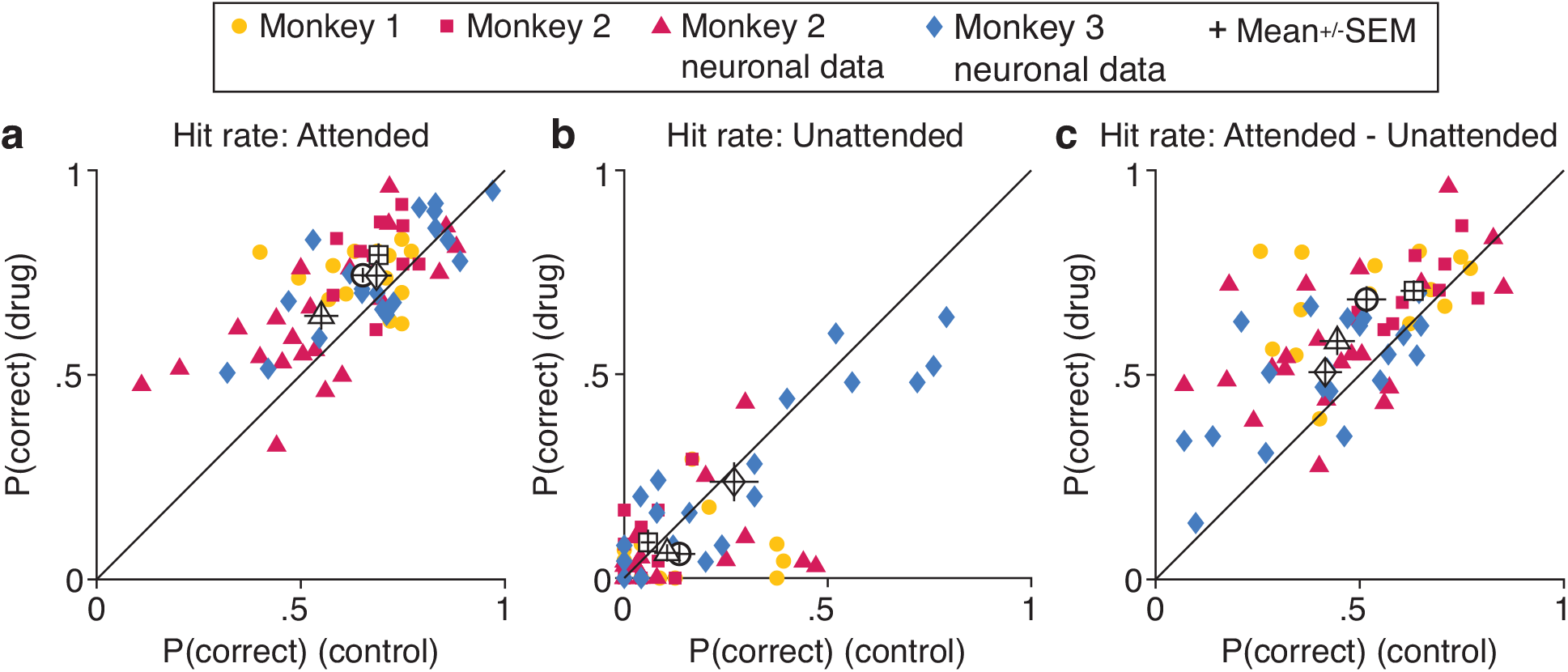
Methylphenidate selectively improves performance at the attended location. (**a**) All three monkeys (marker symbols; see **Methods**) were better able to detect subtle orientation changes at the attended location on drug days (y-axis; numbers represent the hit rate: number of hits divided by hits plus misses) compared to paired control days (x-axis). Attended performance per stimulus location (left or right location; **Fig. 1a**) plotted separately per day. The open symbols and error bars depict the mean and standard error of the mean for each data set. The drug-related improvement was significant for each data set (paired t-tests; Monkey 1: n = 14 [7 pairs of days × 2 stimulus locations per pair], t(13) = -2.5, p = 0.025; Monkey 2: n = 10, t(9) = -3.3, p = 9.2 × 10^-3^; Monkey 2 neuronal dataset: n = 22, t(21) = -3.1, p = 5.6 × 10^-3^; Monkey 3 neuronal dataset: n = 20, t(19) = -2.6, p = 0.019). (**b**) Methylphenidate does not significantly change performance at the unattended location (paired t-tests; Monkey 1: t(13) = 1.8, p = 0.093; Monkey 2: t(9) = -1.0, p = 0.34; Monkey 2 neuronal dataset: t(21) = 1.4, p = 0.17; Monkey 3 neuronal dataset: t(19) = 1.3, p = 0.22). Conventions as in (**a**). (**c**) Comparing the results in (**a**) and (**b**) illustrates that methylphenidate increases the selective effect of attention, defined here as the attention-related difference in hit rate (paired t-tests; Monkey 1: t(13) = -3.5, p = 4.0 × 10^-3^; Monkey 2: t(9) = -2.8, p = 0.019; Monkey 2 neuronal dataset: t(21) = -3.6, p = 1.8 × 10^-3^; Monkey 3 neuronal dataset: t(19) = -2.9, p = 8.5 × 10^-3^). Conventions as in (**a**).

### Spatial specificity in neuronal activity

This spatial specificity in the behavioral effect of methylphenidate was reflected in the V4 neuronal population responses. Consistent with our basic science hypothesis about a general neural coding principle (Ni et al., 2018), methylphenidate improves performance exactly when it changes correlated variability in visual cortex (the average spike count correlation across all simultaneously recorded pairs of V4 neurons; spike count correlation, also called noise correlation, quantifies the trial-to-trial response variability that is shared between a pair of neurons in response to repeated presentations of the same stimulus; Cohen & Kohn, 2011).

Methylphenidate decreased the correlated variability of the recorded V4 neurons only when the animal attended to the stimulus within the receptive fields of the recorded neurons (**Fig. 4a**). It did not decrease the correlated variability when the animal did not attend the stimulus within the neuronal receptive fields (**Fig. 4b**), such that it overall increased the selective effects of attention (the difference in correlated variability between the attended and unattended locations; **Fig. 4c**). These data illustrate a consistent, quantitative relationship between behavioral performance and correlated variability per monkey (**Fig. 4d**), with methylphenidate simply moving the attended behavior and neurons along that quantitative relationship. In other words, the extent to which methylphenidate improved performance at the attended location was matched by the extent to which methylphenidate decreased correlated variability. There was a strong relationship between correlated variability and both visual sensitivity and criterion (**Supp. Fig. 4**; also see Luo & Maunsell, 2015). In contrast, there was no detectable relationship between performance and firing rate for either the drug or placebo control days (**Supp. Fig. 5**).

**Figure 4.**
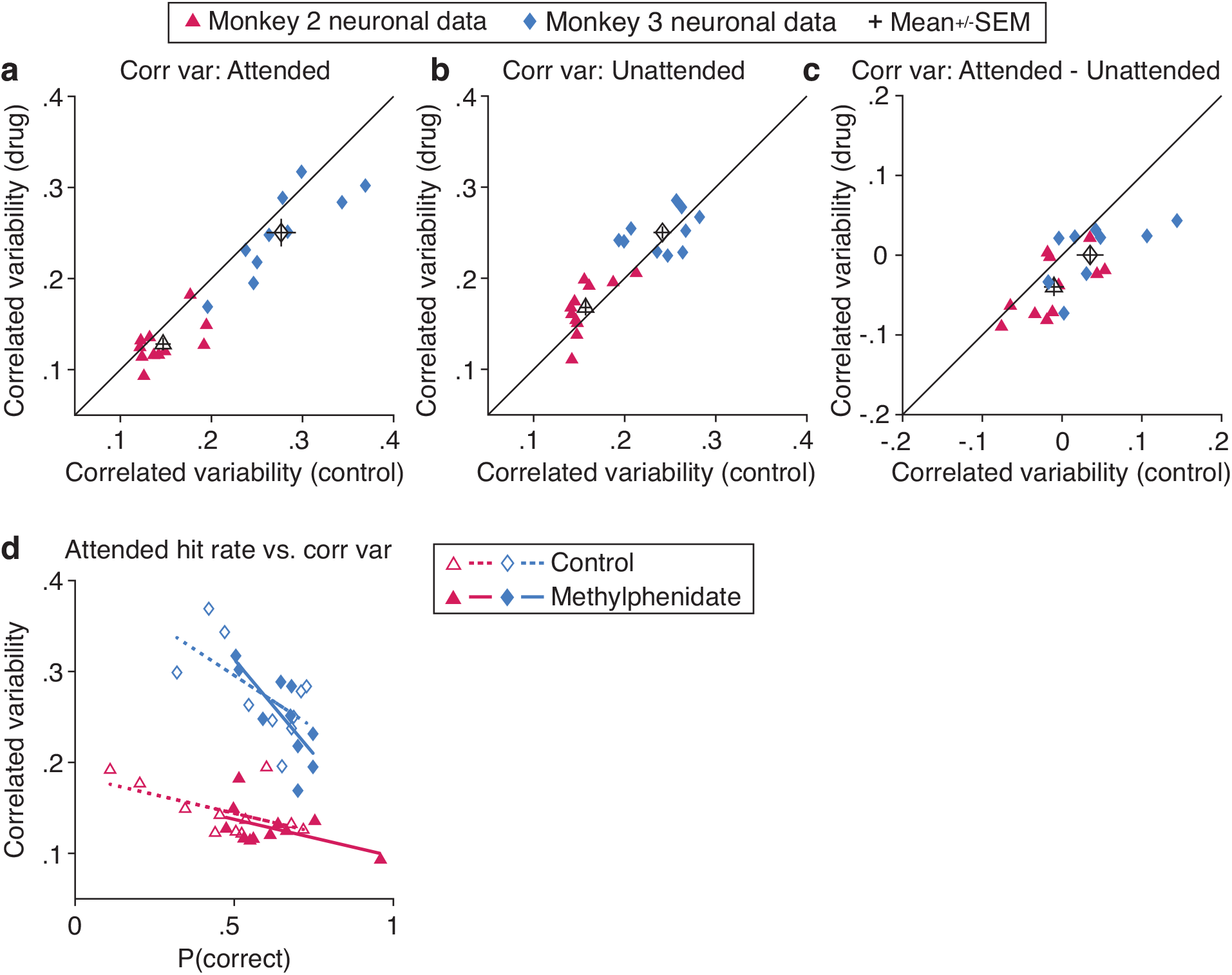
Consistent with our basic science hypothesis, methylphenidate improves performance exactly when it changes correlated variability in visual cortex. (**a**) Methylphenidate reduces V4 correlated variability when the animal pays attention to the joint receptive fields of the recorded neurons. The plot depicts the average noise correlation between all simultaneously recorded neurons on matched drug days (y-axis) and placebo control days (x-axis) for the Monkey 2 and Monkey 3 neuronal datasets (marker symbols; see **Methods**). The mean correlated variability is consistently lower when the receptive field location is attended (paired t-tests; Monkey 2: n = 11 [11 pairs of days x 1 receptive field stimulus location], t(10) = 2.6, p = 0.025; Monkey 3: n = 10, t(9) = 2.9, p = 0.018). The open symbols and error bars depict the mean and standard error of the mean for each data set. (**b**) Methylphenidate does not significantly change V4 correlated variability when the receptive field location is unattended (paired t-tests; Monkey 2: t(10) = -1.7, p = 0.13; Monkey 3: t(9) = -0.89, p = 0.40). Conventions as in (**a**). (**c**) Comparing the results in (**a**) and (**b**) illustrates that methylphenidate increases the selective effect of attention, defined here as the attention-related difference in correlated variability (paired t-tests; Monkey 2: t(10) = 2.9, p = 0.015; Monkey 3: t(9) = 2.7, p = 0.025). (**d**) There is a single, robust relationship between attended behavioral performance (hit rate; x-axis) and attended mean correlated variability (y-axis) for Monkey 2 (correlation coefficient; R = -0.60, p = 3.0 × 10^-3^; correlation was indistinguishable between control and drug conditions, depicted with open and filled symbols, respectively; control: R = -0.55, p = 0.081; drug: R = -0.50, p = 0.11; Fisher z PF test of the difference between dependent but non-overlapping correlation coefficients: zpf = -0.14, p = 0.89) and Monkey 3 (correlation coefficient; R = -0.69, p = 7.9 × 10^-4^; correlation was indistinguishable between control and drug conditions; control: R = -0.63, p = 0.053; drug: R = - 0.76, p = 0.011; Fisher z PF test: zpf = 0.70, p = 0.49). As with natural cognitive processes (control data; also see Ni et al., 2018), systemically administered methylphenidate improves behavioral performance according to the correlated variability change it induces. Best fit lines depicted for control (dashed lines) and methylphenidate data (solid lines).

## DISCUSSION

Cognitive processes like attention can affect performance in a highly selective manner, improving detection of specific stimuli (Maunsell, 2015). This selectivity is often the goal of stimulant use. People use stimulants both with and without prescription with the goal of enhancing selective cognitive processes such as the ability to focus on one task or one aspect of the environment while ignoring distractions (Bagot & Kaminer, 2014; Maher, 2008; Swanson et al., 2011; Wickens et al., 2011). Yet, while we have progressed our understanding of the neuronal mechanisms underlying the effects of these drugs on memory, learning, cognitive flexibility, motivation, and impulsivity (Berridge & Arnsten, 2015; Clatworthy et al., 2009; Devilbiss & Berridge, 2008; Dinse et al., 2003; Dodds et al., 2008; Gamo et al., 2010; Garrett et al., 2015; Kodama et al., 2017; Mehta et al., 2000; Rajala et al., 2012; 2015; 2020), we have only begun to understand the neuronal effects of these stimulants on selective attention in the context of a controlled laboratory setting (Bain et al., 2003; Prendergast et al., 1998; Tomasi et al., 2011; Tremblay et al., 2019). The neural mechanisms underlying stimulant-related changes in selective cognition have remained a mystery: our study is to our knowledge the first electrophysiological report of how changes in neuronal population responses correspond to increased selective attention with ADHD drugs.

Our results demonstrate that a systemic manipulation can selectively change behavior and the underlying neural mechanisms. They support the hypothesis that the spatially selective behavioral and neuronal changes we observed involved an interaction between the diffuse activity of neurotransmitters at the level of top-down control areas (as suggested by in vitro and in vivo measurements of stimulant effects; for review, see Arnsten, 2006; Heal et al., 2013; Mueller et al., 2017) and the localized activity of neurotransmitters at the level of early sensory areas like V4 (as suggested by in vitro and in vivo studies of attention effects; Noudoost & Moore, 2011a; for review, see Deco & Thiele, 2009; Noudoost & Moore, 2011b; Schmitz & Duncan, 2018). While electrophysiological studies have differed in their findings regarding the role of prefrontal cortex in mediating the behavioral effects of methylphenidate (Devilbiss & Berridge, 2008; Gamo et al., 2010; Noudoost & Moore, 2011a; Rajala et al., 2020; Tremblay et al., 2019), the combined global and selective changes we observed here support that global processes can interact with frontoparietal networks (Engelmann et al., 2009; Padmala & Pessoa, 2011) through dopaminergic projections (Botvinick & Braver, 2015; Noudoost & Moore, 2011b) to enhance selective attention processing (Corbetta & Shulman, 2002; Kastner & Ungerleider, 2000; Moore & Zirnsak, 2017; Mueller et al., 2017). Determining how ADHD drugs act through different sites within the brain’s attentional network to enhance selective attention remains an exciting future avenue for both basic and translational neuroscience.

More broadly, our study illustrates that when it comes to combining basic science and translational approaches, the whole is greater than the sum of its parts. We discovered novel behavioral effects of a drug that is widely used, and we leveraged that drug to conduct a strong causal test of a basic science hypothesis that has wide implications for neural coding in many species, systems, and brain areas (Ni et al., 2020; Ruff et al., 2018). Extending this framework to study potential treatments of disorders that affect cognition has the potential to simultaneously transform our understanding of both basic neural mechanisms and clinical outcomes.

## METHODS

The subjects were three adult male rhesus monkeys (*Macaca mulatta*): Monkeys 1, 2, and 3 (7.5, 9.0, and 9.5 kg, respectively). All animal procedures were approved by the Institutional Animal Care and Use Committees of the University of Pittsburgh and Carnegie Mellon University. Each animal was implanted with a titanium head post prior to beginning behavioral training.

### Methylphenidate administration

We tested the behavioral and electrophysiological effects of methylphenidate hydrochloride (Mallinckrodt Pharmaceuticals, St. Louis, MO). Methylphenidate was administered on alternating data collection days (these did not include days on which data were not collected or days on which an insufficient number of trials were collected – see **Data analysis**) for several weeks, providing a minimum of a 24-hour washout period following drug administration prior to collecting control day data (Kodama et al., 2017). A 24-hour washout period between drug and control days was selected based on measurements of orally administered methylphenidate plasma concentrations in rhesus monkeys that determined the drug’s half-life to be less than two hours (Doerge et al., 2000), such that it is undetectable after 12 hours (Oemisch et al., 2016).

On drug administration days, the methylphenidate was dissolved in 10 ml of sugar water (200 mg/ml) and administered orally (the method of dissolving the drug in a flavored liquid for oral administration was adapted from Soto et al., 2012). On control days, 10 ml of sugar water alone (200 mg/ml) was administered orally. For the data in this study, the methylphenidate in sugar water or the sugar water alone was always administered 30 minutes prior to the monkey beginning the change-detection task (based on prior studies that used similar rhesus monkey behavioral session timing after oral stimulant administration; Gamo et al., 2010; Rajala et al., 2012; 2015).

A maximum dosage of 8.0 mg/kg was pre-determined based on prior studies performed in rhesus monkeys (Czoty et al., 2013; Gamo et al., 2010; Rajala et al., 2012; 2015; Soto et al., 2012). The dosages included in the analyses were 2.0, 3.0, 4.0, 5.0, and 6.0 mg/kg (**Supp. Fig. 2a, b**). Dosages of 6.0 and 7.0 mg/kg sometimes led to agitation that prevented the monkeys from being able to perform the task. This occurred with 1 out of 1 test of 6.0 mg/kg for Monkey 1, 1 out of 2 tests of 6.0 mg/kg for Monkey 2, and 1 out of 1 test of 7.0 mg/kg for Monkey 2. Due to these effects, we did not test higher than 5.0 mg/kg with Monkey 3, and we never tested a dosage higher than 7.0 mg/kg. The mean analyzed dosage was 3.8 mg/kg (doses of 3.0 mg/kg in rhesus macaques result in similar plasma levels as therapeutic doses of 0.3 mg/kg in humans; Doerge et al., 2000).

Agitation or drowsiness leading to the inability to collect behavioral data has been previously reported at higher stimulant dosages (Rajala et al., 2012; Kodama et al., 2017). Here, the agitating effect of higher dosages described above manifested as an increase in erratic eye movements, resulting in an inability to fixate and initiate behavioral trials. This decrease in stimulant efficacy at higher dosages follows the characteristic inverted U-shaped pharmacological dose-response curve (Calabrese & Baldwin, 2001) that has been well documented for stimulants (Borota et al., 2014; Dodds et al., 2008; Gamo et al., 2010; Martelle et al., 2013; Rajala et al., 2012; for review, see Fredholm et al., 1999; Noudoost & Moore, 2011b; Swanson et al., 2011).

Data from all dosages were combined for each analysis to avoid best dose analysis (Soto et al., 2013), as our goal was to use methylphenidate as a causal mechanism to test our hypotheses, not to test for dose-dependent effects (see Rajala et al., 2012 for analyses of methylphenidate dose-dependent effects in rhesus monkeys).

### Behavioral task

The monkeys performed an orientation change-detection task (Cohen & Maunsell, 2009; Ni et al., 2018) with cued attention (Posner, 1980). All three monkeys were trained extensively on this task before the data presented here were recorded. Visual stimuli were presented on a CRT monitor (calibrated to linearize intensity; 1024 × 768 pixels; 120 Hz refresh rate) placed 57 cm from the monkey, using custom software written in MATLAB (Psychophysics Toolbox; Brainard, 1997; Pelli, 1997). Eye position was monitored using an infrared eye tracker (Eyelink 1000; SR Research) as per previously published methods (Ni et al., 2018).

A monkey began a trial by fixing its gaze on a small spot presented in the center of the video display (**Fig. 1a**). Next, two peripheral drifting Gabor stimuli, one presented in the left visual hemifield and one presented in the right visual hemifield, synchronously flashed on (for 200 ms) and off (for an interval that was randomly selected from a uniform distribution with a range of 200-400 ms) until, at a random and unsignaled time, the orientation of one of the stimuli changed. The monkey received a liquid reward for making a saccade to the changed stimulus within 450 ms of its onset and was randomly administered extra rewards after correctly completed trials. If no orientation change occurred within a maximum of 12-15 stimulus presentations (∼10% of the trials), the trial was terminated and the monkey received a liquid reward simply for having maintained fixation throughout the trial (catch trials).

The size, two locations, temporal frequency, and spatial frequency of the Gabor stimuli were fixed for both days of a pair (the drug day and the paired placebo control day). The orientation change amount was also fixed for both days of a pair, and was the same for both stimulus locations and all trials. The starting orientation at which each stimulus was flashed multiple times before any orientation change occurred was selected randomly per trial and per stimulus location from a set of 4-12 different starting orientations.

The attended location alternated between the left and right stimulus locations (**Fig. 1a**) on each new block of 120-125 trials. Prior to a new block, the monkey was cued to attend to one stimulus location with 10 instruction trials in which a stimulus was only flashed at that one location. During each block, the orientation change occurred at the cued location on 80% of the trials and at the other location on 20% of the trials.

### Data sets

During the behavioral data sets (collected for Monkey 1 and Monkey 2 and illustrated with circle markers and square markers, respectively), no neuronal data were collected. The monkey controlled the length of each experimental session: the session ended when the monkey had not fixated the central fixation point to initiate a trial for 10 minutes. For each monkey, the two locations for the Gabor stimuli were selected based on the monkey demonstrating approximately equal performance at those two locations prior to beginning data collection.

During the neuronal data sets (collected for Monkey 2 and Monkey 3 and illustrated with triangle markers and diamond markers, respectively), psychophysical and neuronal data were collected simultaneously. For each monkey, the two locations for the Gabor stimuli were selected such that one location maximally overlapped the joint recorded receptive fields and the other location was in the opposite visual hemifield.

### Neurophysiological recordings

For the neuronal data sets collected for Monkey 2 and Monkey 3, we recorded extracellularly per monkey using a single chronically implanted microarray (48 electrodes per array, Blackrock Microsystems) in visual area V4 (left hemisphere for Monkey 2 and right hemisphere for Monkey 3; each monkey also had a second chronically implanted microarray, the data from which are not included in this study), using previously published methods (Ni et al., 2018). We set the same spike-detection voltage threshold across all electrodes and all recording sessions and included all threshold crossings as the neuronal activity per electrode (the recorded “unit”; Ni et al., 2018; Trautmann et al., 2019; see **Data analysis**). The typical receptive field size plotted in **Fig. 1b** (dotted yellow circle) was calculated as the standard deviation of a Gaussian fit.

### Data analysis

Statistical details can be found in the figure legends (statistical tests used, *n* values, etc.). Experimental sessions were included in the analyses if a minimum of 200 change-detection trials were completed (correct or incorrect).

To determine the effect of methylphenidate on the amount of time a monkey engaged in the change-detection task (**Fig. 2, Supp. Fig. 1, Supp. Fig. 2c, d**), the behavioral data sets were analyzed. The time engaged in the task was calculated as the time between the start time of the first trial and the end time of the tenth from last correctly completed trial (excluding the last trials conservatively estimated the working time so as to not include potential breaks between periods of concerted effort near the end of the session). The results were qualitatively unchanged when the total experimental time (from the start time of the first trial to the end time of the 10 minute break that ended the session) was analyzed instead (paired *t*-tests; Monkey 1: *n* = 7 pairs of days, *t*(6) = -4.2, *p* = 5.7 × 10^-3^; Monkey 2: *n* = 5 pairs of days, *t*(4) = -3.8, *p* = 0.019). To determine the effect of methylphenidate on performance (**Fig. 3, Fig. 4c, Supp. Fig. 2**-**5**), the behavioral and/or neuronal data sets were analyzed. For analyses of performance, only the first two blocks collected per experimental session were analyzed (one block with attention cued to the left hemifield stimulus location, one block with attention cued to the right hemifield stimulus location; **Fig. 1**). Only the first two blocks were analyzed per experimental session to control for potential changes in drug efficacy and motivation levels across the session. Instruction and catch trials were not included in the analyses.

To determine the effect of methylphenidate on neuronal population activity (**Fig. 4, Supp. Fig. 4-5**), the neuronal data sets were analyzed. Recorded units were included in the analyses on a pair-by-pair basis. The same units were analyzed for both days of a pair, based on the responses of the units on the control day of the pair: the analyzed units were the units that passed a mean stimulus-evoked firing rate of at least 10 Hz and a mean stimulus-evoked firing rate that was significantly higher than the mean firing rate during a baseline period in which no stimuli were presented (stimulus analysis period: 60-200 ms from stimulus onset to account for V4 response latency; baseline analysis period: 100 ms interval prior to the onset of the first stimulus/trial; included trials: completed orientation-change and catch trials; included stimuli: all stimuli but the first stimulus/trial and any orientation-change stimuli; based on a two-sided Wilcoxon signed rank test of whether the response ratio of the mean stimulus-evoked firing rate compared to the mean baseline firing rate was different from 1). Results were not qualitatively different when these same criteria were applied on a day-by-day basis (applied to each session individually, regardless of day pairing). The population size of simultaneously recorded units included in the analyses was 26-32 units for Monkey 2 (mean 30) and 3-29 units for Monkey 3 (mean 17).

To analyze the firing rates and correlated variability of the V4 neuronal populations in response to stimuli presented at the receptive field location (**Fig. 1b**), stimuli presented during attended orientation-change, catch, and false alarm trials (the attended condition) were compared to stimuli presented during unattended orientation-change, catch, and false alarm trials (the unattended condition). All stimuli were included except the first stimulus per trial, orientation-change stimuli, and stimulus presentations during which the monkey made a false alarm (a saccade to a stimulus location where no orientation change had occurred). The neuronal responses to a stimulus were calculated during the analysis period of 60-260 ms from stimulus onset.

The neuronal population correlated variability was calculated as the mean (across all pairs of units) correlation coefficient between the responses of two units to repeated presentations of the same stimulus. The correlation coefficient per pair of units was calculated per starting orientation and averaged across all starting orientations. Correlation coefficients >0.5 and <-0.1 were excluded from mean calculations.

## Data availability

Electrophysiological data analyzed in this manuscript will be available at https://github.com/amymni/.

## Code availability

Any original code used for this manuscript will be available at https://github.com/amymni/.

## ACKNOWLEDGEMENTS

A.M.N. received support from US NIH grant 1K99NS118117-01 and a fellowship from the Simons Foundation Collaboration on the Global Brain. M.R.C. received support from US NIH grants 4R00EY020844-03, R01 EY022930, R01NS121913, and Core Grant P30 EY008098s, a grant from the Whitehall Foundation, a Klingenstein-Simons Fellowship, a Sloan Research Fellowship, a McKnight Scholar Award, and a grant from the Simons Foundation Collaboration on the Global Brain. We thank Nancy Ator for guidance and discussions throughout this study. We thank Lily E. Kramer for assistance with data collection. We thank Karen McCracken for technical assistance. We thank John H.R. Maunsell, Julio C. Martínez-Trujillo, Sébastien Tremblay, and Cheng Xue for comments on previous versions of this manuscript.

## AUTHOR CONTRIBUTIONS

A.M.N., B.S.B., D.A.R., and M.R.C. designed the experiment. A.M.N., B.S.B., and D.A.R. collected the data. A.M.N. performed the analyses. A.M.N., D.A.R., and M.R.C. wrote the paper.

## SUPPLEMENTARY FIGURES

**Supplementary Figure 1.**
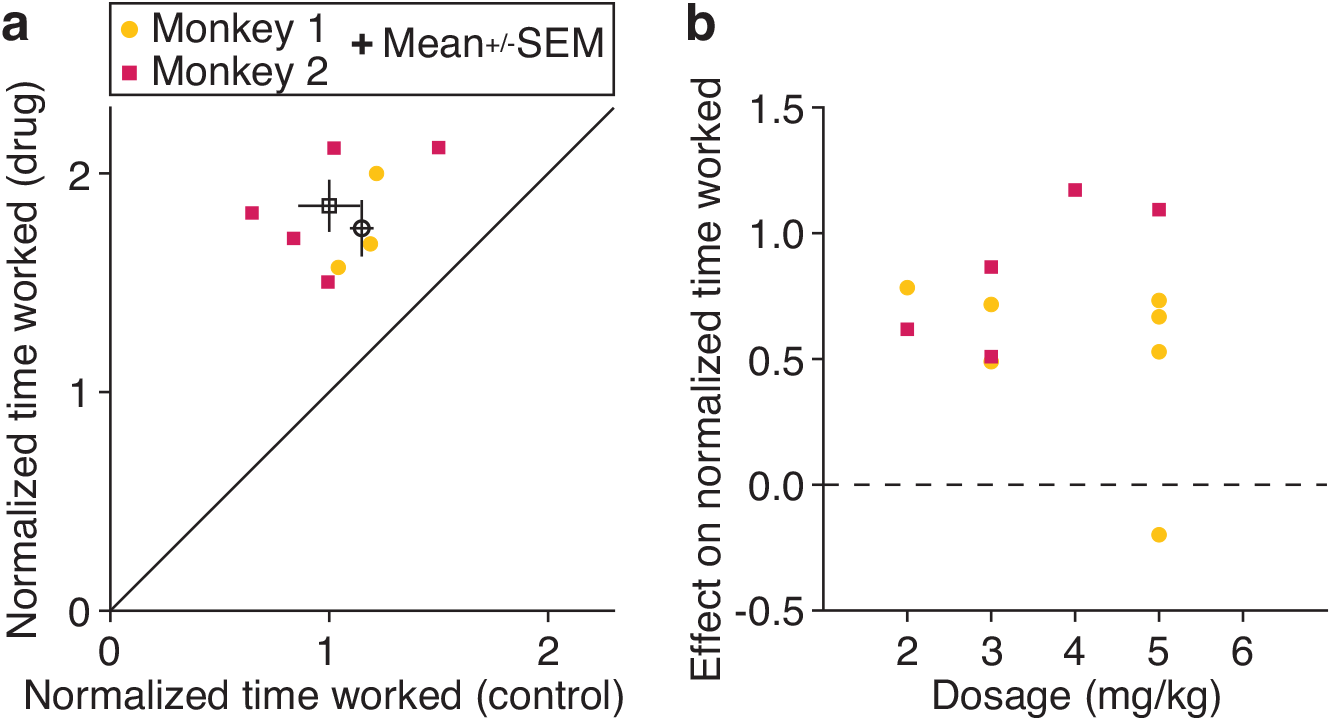
The effect of methylphenidate on working time does not depend on water consumption or methylphenidate dosage. (**a**) The plot depicts the amount of time the monkey engaged in the change-detection task, normalized to the mean time worked on placebo control days. Each point is the normalized working time for a matched drug day (*y*-axis) and control day (*x*-axis) for each monkey (marker symbols). The open symbols are the mean for each monkey, and error bars represent standard error of the mean (SEM). We subsampled our data so that the mean liquid consumption was indistinguishable before drug and control days for each monkey. In this subset of data, the significant methylphenidate-related increase in working time persists (paired *t*-tests; Monkey 1: *n* = 3 pairs of days, *t*(2) = -6.5, *p* = 0.023; for Monkey 2 mean liquid consumption was already indistinguishable before drug and control days and thus the data match the data in the main text: *n* = 5 pairs of days, *t*(4) = -6.6, *p* = 2.7 × 10^-3^). (**b**) The effect of methylphenidate on the time the monkey engaged in the change-detection task (*y*-axis; normalized time engaged on the drug day – normalized time engaged on the matched control day) is not consistently related to methylphenidate dosage (*x*-axis; Kendall’s rank correlation coefficient; Monkey 1: *n* = 7 pairs of days, *τ* = -0.17, *p* = 0.49; Monkey 2: *n* = 5 pairs of days, *τ* = 0.60, *p* = 0.031; though see Rajala et al., 2012).

**Supplementary Figure 2.**
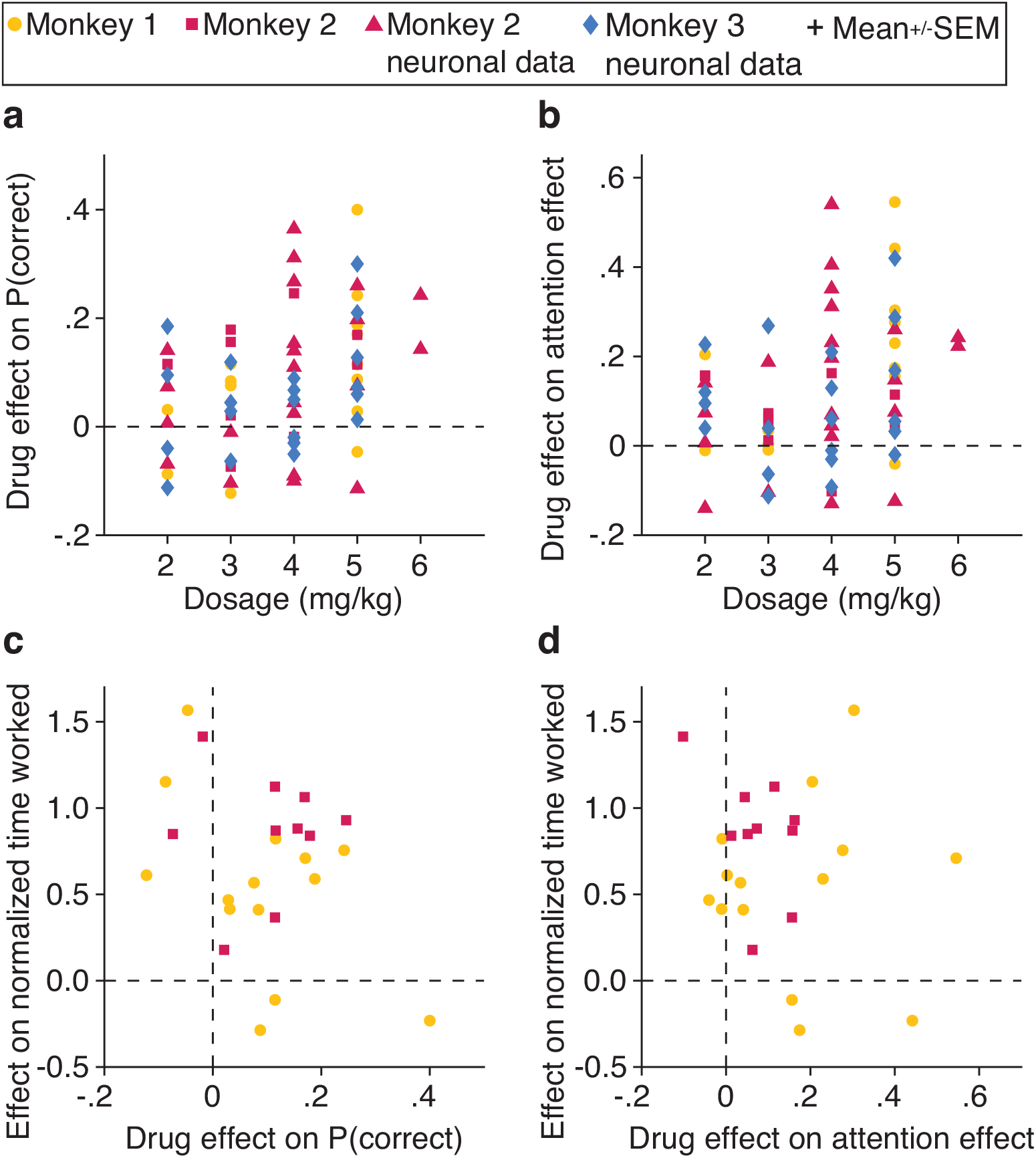
The effect of methylphenidate on performance does not depend on dosage or on the effect of methylphenidate on working time. (**a**) The effect of methylphenidate on performance at the attended location (*y*-axis; attended hit rate on the drug day – attended hit rate on the paired control day) is not significantly related to methylphenidate dosage (*x*-axis) for each data set (marker symbols; Kendall’s rank correlation coefficient; Monkey 1: *n* = 14 [7 pairs of days × 2 stimulus locations per pair], *τ* = 0.45, *p* = 0.054; Monkey 2: *n* = 10, *τ* = 0.15, *p* = 0.64; Monkey 2 neuronal dataset: *n* = 22, *τ* = 0.24, *p* = 0.16; Monkey 3 neuronal dataset: *n* = 20, *τ* = 0.27, *p* = 0.13). (**b**) The effect of methylphenidate on selective attention (*y*-axis; the difference in hit rate between the attended and unattended locations on the drug day – the difference in hit rate between the attended and unattended locations on the paired control day) is not significantly related to methylphenidate dosage (*x*-axis; Kendall’s rank correlation coefficient; Monkey 1: *τ* = 0.45, *p* = 0.054; Monkey 2: *τ* = -0.25, *p* = 0.40; Monkey 2 neuronal dataset: *τ* = 0.25, *p* = 0.14; Monkey 3 neuronal dataset: *τ* = 0.072, *p* = 0.71). (**c**) There is no detectable relationship between the effect of methylphenidate on performance at the attended location (*x*-axis; attended hit rate at one stimulus location on the drug day – attended hit rate at the same stimulus location on the paired control day) and the effect of methylphenidate on the time the monkey engaged in the change-detection task (*y*-axis; normalized time engaged at one stimulus location on the drug day – normalized time engaged at the same stimulus location on the matched control day) for each monkey (correlation coefficient; Monkey 1: *R* = -0.50, *p* = 0.069; Monkey 2: *R* = 0.035, *p* = 0.92). Time worked is normalized to the mean time worked on the placebo controls of the pairs. (**d**) There is no detectable relationship between the effect of methylphenidate on selective attention (*x*-axis; the difference in hit rate between attending and not attending one stimulus location on the drug day – the difference in hit rate between attending and not attending the same stimulus location on the paired control day) and the effect of methylphenidate on the time the monkey engaged in the change-detection task (*y*-axis; normalized time engaged at one stimulus location on the drug day – normalized time engaged at the same stimulus location on the matched control day) for each monkey (correlation coefficient; Monkey 1: *R* = 0.027, *p* = 0.93; Monkey 2: *R* = -0.45, *p* = 0.19). It should be noted that it was not our goal to test for dose-dependent effects, and that prior studies have found that the same stimulant can have different effects on different cognitive processes depending on the dosage administered (Pietrzak et al., 2006; Rajala et al., 2012; 2020; Swanson et al., 2011; Wickens et al., 2011).

**Supplementary Figure 3.**
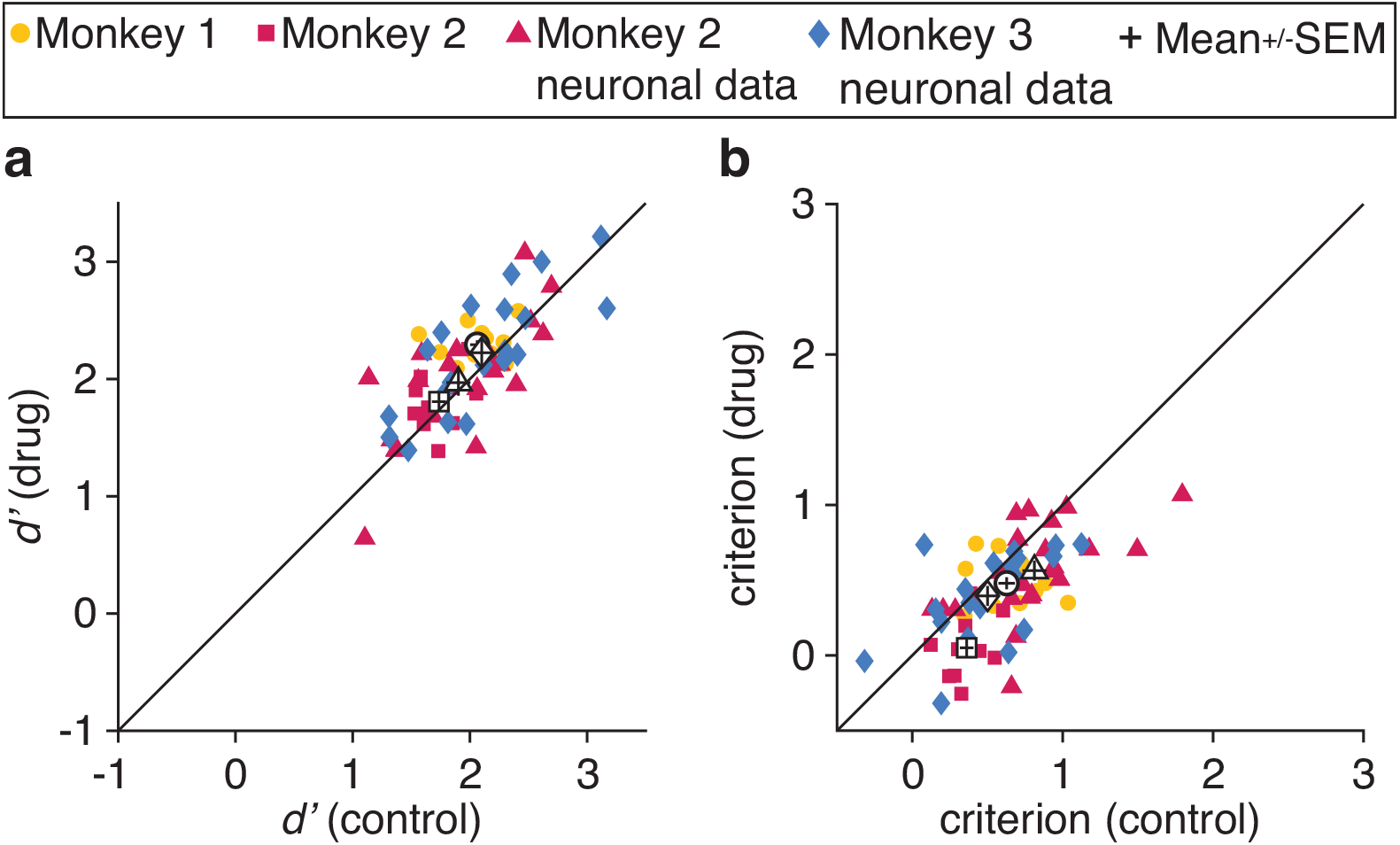
Methylphenidate increases hit rate at the attended location by both increasing visual sensitivity and decreasing criterion. (**a**) Methylphenidate improved sensitivity (*d’*) at the attended location on drug days *(y*-axis) compared to paired control days (*x*-axis) across the entire data set (paired *t*-test: *t*(65) = -3.0, *p* = 3.4 × 10^-3^), though not significantly for all individual data sets (paired *t*-tests; Monkey 1: *n* = 14 [7 pairs of days × 2 stimulus locations per pair], *t*(13) = -3.4, *p* = 4.7 × 10^-3^; Monkey 2: *n* = 10, *t*(9) = -0.87, *p* = 0.41; Monkey 2 neuronal dataset: *n* = 22, *t*(21) = -0.87, *p* = 0.40; Monkey 3 neuronal dataset: *n* = 20, *t*(19) = -1.6, *p* = 0.12). The open symbols and error bars depict the mean and standard error of the mean for each data set (marker symbols). (**b**) Methylphenidate decreased criterion at the attended location on drug days compared to paired control days across the entire data set (paired *t*-test: *t*(65) = 5.3, *p* = 1.3 × 10^-6^) though not significantly for all individual data sets (paired *t*-tests; Monkey 1: *t*(13) = 2.1, *p* = 0.059; Monkey 2: *t*(9) = 4.8, *p* = 9.2 × 10^-4^; Monkey 2 neuronal dataset: *t*(21) = 3.6, *p* = 1.8 × 10^-3^; Monkey 3 neuronal dataset: *t*(19) = 1.6, *p* = 0.13). Conventions as in (**a**). It is not surprising that methylphenidate affects both sensitivity and criterion because these measures have been demonstrated to be strongly yoked (Luo & Maunsell, 2018; Sridharan et al., 2017). Attentional measures that improve performance generally affect both sensitivity and criterion (Luo & Maunsell, 2015).

**Supplementary Figure 4.**
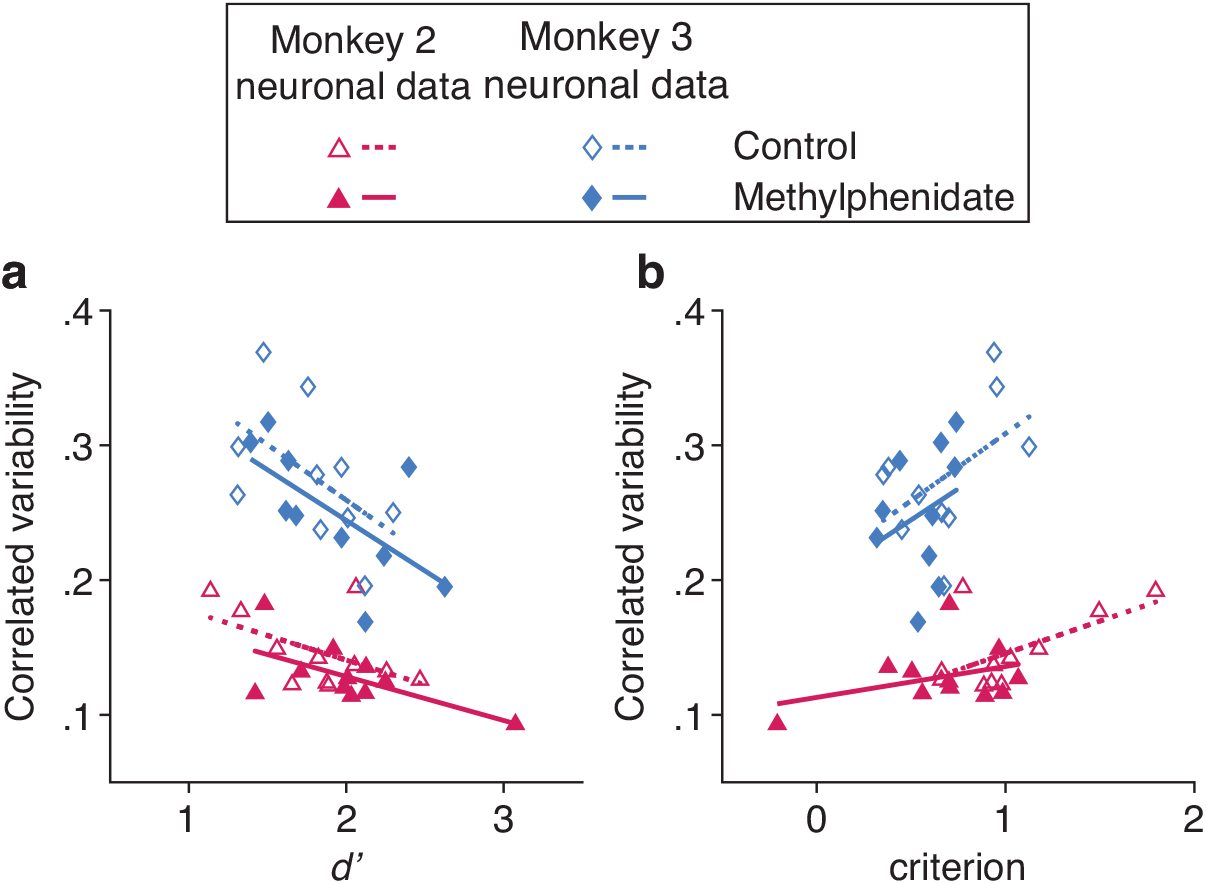
Methylphenidate both improves visual sensitivity and decreases criterion when it changes correlated variability in V4. (**a**) There was a single relationship between visual sensitivity at the attended location *(d’*; *x*-axis) and attended mean correlated variability (*y*-axis) for Monkey 2 (correlation coefficient; *R* = -0.59, *p* = 3.8 × 10^-3^; correlation was indistinguishable between control and drug conditions, depicted with open and filled symbols, respectively; control: *n* = 11 days, *R* = -0.51, *p* = 0.11; drug: *n* = 11 days, *R* = -0.63, *p* = 0.038; Fisher *z* PF test of the difference between dependent but non-overlapping correlation coefficients: *zpf* = 0.40, *p* = 0.69) and Monkey 3 (correlation coefficient; *R* = -0.61, *p* = 4.4 × 10^-3^; control: *n* = 10 days, *R* = -0.54, *p* = 0.11; drug: *n* = 10 days, *R* = -0.65, *p* = 0.043; Fisher *z* PF test: *zpf* = 0.40, *p* = 0.69). Best fit lines depicted for control (dashed lines) and methylphenidate data (solid lines). (**b**) There was a single relationship between criterion at the attended location *(x*-axis) and attended mean correlated variability (*y*-axis) for Monkey 2 (correlation coefficient; *R* = 0.57, *p* = 5.4 × 10^-3^; control: *R* = 0.60, *p* = 0.051; drug: *R* = 0.36, *p* = 0.28; Fisher *z* PF test: *zpf* = 0.72, *p* = 0.47) and Monkey 3 (correlation coefficient; *R* = 0.46, *p* = 0.041; control: *R* = 0.51, *p* = 0.13; drug: *R* = 0.20, *p* = 0.42; Fisher *z* PF test: *zpf* = 0.89, *p* = 0.37). Conventions as in (**a**).

**Supplementary Figure 5.**
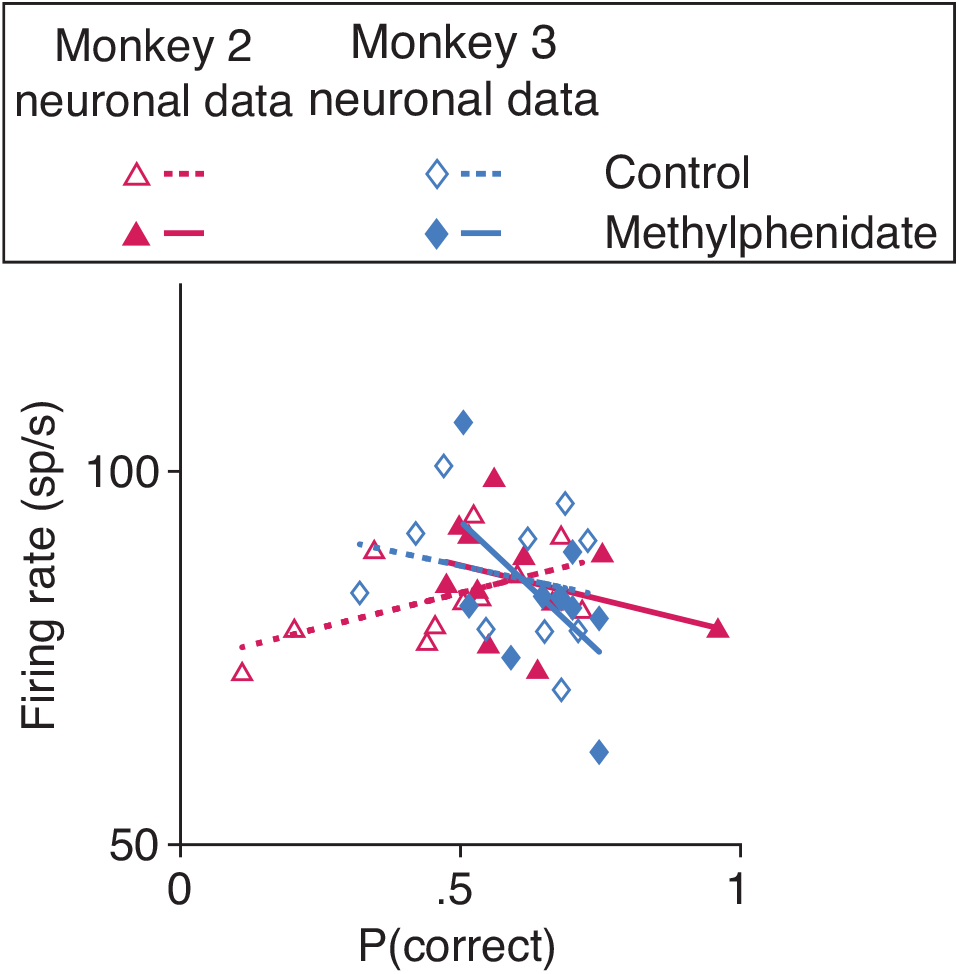
Unlike with correlated variability, there was no detectable relationship between performance at the attended location (hit rate; *x*-axis) and attended mean firing rate (*y*-axis) for Monkey 2 (correlation coefficient; *R* = 0.18, *p* = 0.42; control and drug conditions depicted with open and filled symbols, respectively; control: *n* = 11 days, *R* = 0.54, *p* = 0.084; drug: *n* = 11 days, *R* = -0.34, *p* = 0.30) or for Monkey 3 (correlation coefficient; *R* = -0.39, *p* = 0.093; control: *n* = 10 days, *R* = -0.24, *p* = 0.51; drug: *n* = 10 days, *R* = -0.56, *p* = 0.093). Best fit lines depicted for control (dashed lines) and methylphenidate data (solid lines).

